# Attenuation of the extracellular matrix increases the number of synapses but suppresses synaptic plasticity

**DOI:** 10.1101/2020.04.23.058115

**Authors:** Yulia Dembitskaya, Nikolay Gavrilov, Igor Kraev, Maxim Doronin, Olga Tyurikova, Alexey Semyanov

**Affiliations:** Department of Molecular Neurobiology, Shemyakin-Ovchinnikov Institute of Bioorganic Chemistry, Russian Academy of Sciences, Miklukho-Maklaya street 16/10, Moscow, 117997, Russia; Lobachevsky State University of Nizhny Novgorod, Nizhny Novgorod, 603950, Russia; Electron Microscopy Suite, Faculty of Science, Technology, Engineering and Mathematics, Open University, Milton Keynes MK7 6AA, UK; Sechenov First Moscow State Medical University, Bolshaya Pirogovskaya str 19с1, Moscow, Russia, 119146

## Abstract

The brain extracellular matrix (ECM) is a proteoglycan complex that occupies the extracellular space between brain cells and regulates brain development, brain wiring, and synaptic plasticity. However, the action of the ECM on synaptic plasticity remains controversial. Here, we employed serial section electron microscopy to show that enzymatic attenuation of ECM with chondroitinase ABC (ChABC) triggers the appearance of new glutamatergic synapses onto thin dendritic spines of CA1 pyramidal neurons. The appearance of new synapses increased the ratio of the field excitatory postsynaptic potential (fEPSP) to presynaptic fiber volley (PrV), suggesting that these new synapses are formed on existing axonal fibers. However, both the mean miniature excitatory postsynaptic current (mEPSC) amplitude and AMPA/NMDA ratio were decreased, suggesting that ECM attenuation increased the proportion of ‘unpotentiated’ synapses. A higher proportion of unpotentiated synapses would be expected to promote long-term potentiation (LTP). Surprisingly, theta-burst induced LTP was suppressed by ChABC treatment. The suppression of LTP was accompanied by decreased excitability of CA1 pyramidal neurons due to the upregulation of small conductance Ca^2+^-activated K^+^ (SK) channels. A pharmacological blockade of SK channels restored cell excitability and, expectedly, enhanced LTP above the level of control. This enhancement of LTP was abolished by a blockade of Rho-associated protein kinase (ROCK), which is involved in the maturation of dendritic spines. Thus, ECM attenuation enables the appearance of new synapses in the hippocampus, which is compensated for by a reduction in the excitability of postsynaptic neurons, thereby preventing network overexcitation at the expense of synaptic plasticity.

---

The components of the extracellular matrix (ECM) form a molecular meshwork in the extracellular space of the brain. The major components of the ECM are hyaluronic acid, chondroitin sulfate proteoglycans (CSPGs), link proteins such as hyaluronan and proteoglycan link protein 1 (HAPLN1/CRTL1), and glycoproteins such as tenascin-R, which cross-link CSPGs and stabilize ECM structure (Morawski et al., 2014). The ubiquitous localization of ECM allows it to broadly affect neuronal function, both through mechanical and electrical regulation of diffusion in the extracellular space, as well as by interacting with several transmembrane receptors and channels (Nicholson and Hrabetova, 2017).

During prenatal and early postnatal brain development, the ECM provides either adhesive or repellent cues that control cell migration and navigation of growing axons. These processes involve signaling through a diverse group of transmembrane receptors, such as integrins (Wehrle-Haller and Bastmeyer, 2014), ApoER2, VLDL, EphB (Bouche et al., 2013; Sharaf et al., 2013), and RPTPσ (Coles et al., 2011; Lang et al., 2015). In the mature brain, a number of ECM molecules and ECM receptors trigger various biochemical cascades that regulate neuronal function (Kerrisk et al., 2014). Attenuation of the ECM in the adult brain can potentially restore the brain to an immature state and promote (re-)wiring (Bikbaev et al., 2015). Hence, manipulation of the ECM has been suggested as a way to stimulate brain repair after injury and boost brain plasticity (Burnside and Bradbury, 2014; Chao et al., 2018). Nevertheless, the role of ECM in synaptic plasticity remains controversial (Dityatev and Schachner, 2003; Senkov et al., 2014). Initial studies suggested that acute enzymatic digestion of CSPGs with chondroitinase ABC (ChABC) impaired both long-term potentiation (LTP) and depression (LTD) in the CA1 region of the hippocampus (Bukalo et al., 2001). Similarly, LTP in hippocampal CA1 neurons was reduced by a deficiency in the ECM components brevican or tenascin-R reduced (Brakebusch et al., 2002; Bukalo et al., 2001, 2007; Saghatelyan et al., 2001), as well as after acute enzymatic digestion of hyaluronic acid with hyaluronidase (Kochlamazashvili et al., 2010). Moreover, intra-hippocampal injection of hyaluronidase impaired contextual fear conditioning (Kochlamazashvili et al., 2010). These studies demonstrate that ECM attenuation may disrupt synaptic plasticity and some forms of learning and memory. On the other hand, mice deficient in CSPG phosphacan/RPTPbeta exhibit increased LTP (Niisato et al., 2005). HAPLN1 knockout mice retain high juvenile-type levels of ocular dominance plasticity in adulthood (Carulli et al., 2010) and exhibit enhanced long-term object recognition memory and LTD in the perirhinal cortex (Romberg et al., 2013). Similarly, recognition memory is enhanced following ChABC injection (Romberg et al., 2013). These findings suggest that genetic or enzymatic attenuation of the ECM may promote some forms of synaptic plasticity and learning.

Here, we resolve this controversy by showing that ECM attenuation with ChABC increases the number of ‘plastic’ glutamatergic synapses onto CA1 pyramidal neurons while at the same time reducing neuronal excitability through the upregulation of small conductance Ca^2+^-activated K^+^ (SK) channels that suppress LTP.

## Results

### ECM attenuation increases the number of glutamatergic synapses onto CA1 pyramidal neurons

Hippocampal slices from two mice (two slices per mouse) were treated with either ChABC or a sham solution and were used for serial section electron microscopy. Dendritic spines and axonal boutons were visually identified, manually segmented, and reconstructed in 3D (Fig. 1a,b). To avoid selection bias, the number of spines and boutons was calculated in five randomly selected block series of each 3D reconstruction (Fig. 1c,d; Fig. S1). The spines were divided into two groups: mushroom spines and thin spines. Mushroom spines, which are also known as ‘memory spines,’ are mature spines and reflect potentiated synapses (Bourne and Harris, 2007). Their density was not significantly affected by ChABC treatment (sham: 0.51 ± 0.05 spines/µm^3^, *n* = 10; ChABC: 0.48 ± 0.05 spines/µm^3^, *n* = 10; *p* = 0.7, two-sample *t*-test; Fig. 1e). Thin spines have been dubbed ‘learning spines’ as they correspond to synapses that can be further potentiated (Bourne and Harris, 2007). ChABC treatment significantly increased the density of thin spines (sham: 1.7 ± 0.1 spines/µm^3^, *n* = 10; ChABC: 3.0 ± 0.2 spines/µm^3^, *n* = 10; *p* < 0.001, two-sample *t*-test; Fig. 1f). In addition, we observed increased density of axonal boutons in the ChABC treated slices (sham: 2.3 ± 0.1 boutons/µm^3^, *n* = 10; ChABC: 3.4 ± 0.3 boutons/µm^3^, *n* = 10; *p* < 0.001, two-sample *t*-test; Fig. 1g). These findings are consistent with an increase in the number of CA1 – CA3 synaptic connections following ECM attenuation.

**Figure 1.**
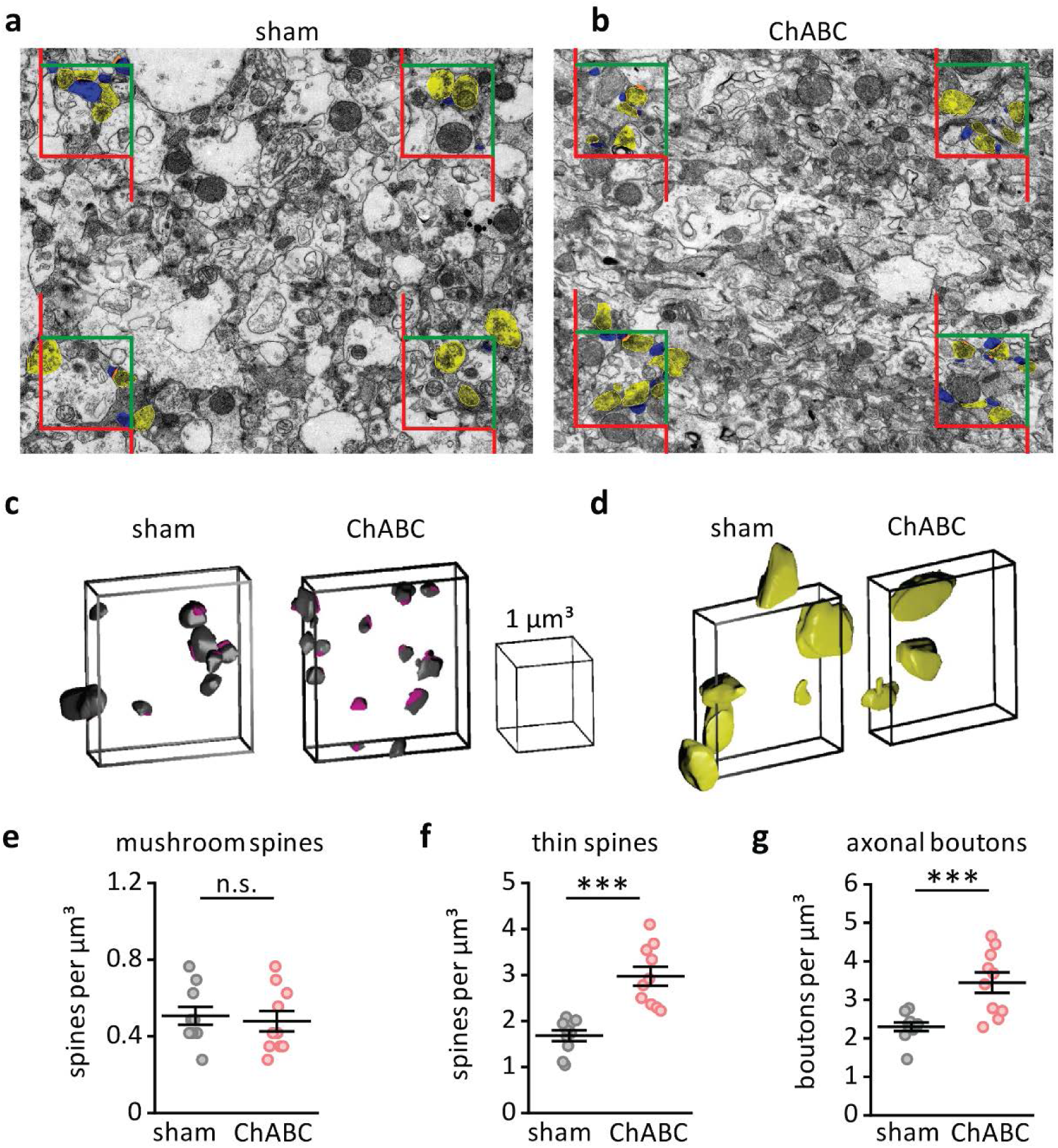
ECM attenuation increases the number of glutamatergic synapses onto CA1 pyramidal neurons. **a,b.** Electron microscopy images from sham-treated (*a*) and ChABC-treated (*b*) slices showing dendritic spines (*blue*), axonal boutons (*yellow*) and postsynaptic densities (*orange*) within and at the borders of unbiased bricks. Green sides of the bricks are inclusive, red sides are exclusive (see Methods section). **c,d.** Examples of 3D reconstructed dendritic spines (*c*) and axonal boutons (*d*) analyzed in the unbiased bricks within the samples. *Left box* - sham, *right box* – ChABC. **e,f,g.** Summary plot of mushroom spine (*e*), thin spine (*f*), and axonal bouton (*g*) densities in 10 boxes from 2 sham-treated slices (*black circles*) and 2 slices treated with ChABC (*pink circles*). The data are presented as the mean ± SEM. n.s. *p* > 0.05; *** *p* < 0.001; two-tailed two-sample *t*-test

### New synapses are formed by existing axons

New synapses can either be associated with the outgrowth of new axons or can be formed from existing fibers. To accurately determine the source of new synapses, we recorded the presynaptic fiber volley (PrV) and field (f) excitatory postsynaptic potential (EPSP) that occurred in response to extracellular stimulation of Schaffer collaterals (Fig. 2a,b). We varied the stimulus intensity and determined the relationship between the fEPSP slope and the PrV amplitude (the input-output curve). The input-output curve significantly shifted upward after ChABC treatment (*F*(1,80) = 17.23, *p* < 0.001, two-way repeated measures [RM] ANOVA; Table S1, Fig. 2c). This finding suggests that the activation of an identical number of fibers produced larger EPSPs in the postsynaptic neuron. This could be a result of existing axons making additional synapses or of the existing synapses becoming potentiated. To distinguish between these two possibilities, we recorded the miniature (m) excitatory postsynaptic currents (EPSCs) in CA1 pyramidal neurons (Fig. 2d). ChABC treatment did not significantly change the frequency of mEPSCs (sham: 0.76 ± 0.07 Hz, *n* = 7; ChABC: 0.61 ± 0.05 Hz, *n* = 7; *p* = 0.07, two-sample *t*-test; Fig. 2e); however, the mean mEPSC amplitude decreased (sham: 22.33 ± 2.20 pA, *n* = 7X; ChABC: 15.50 ± 1.90 pA, *n* = 7; *p* = 0.018, two-sample *t*-test; Fig. 2f), suggesting that ECM attenuation does not potentiate existing glutamatergic synapses onto CA1 pyramidal neurons. Thus, the upward shift of the input-output curve may be explained by an increase in the number of synaptic contacts made by Schaffer collaterals onto CA1 pyramidal neurons.

**Figure 2.**
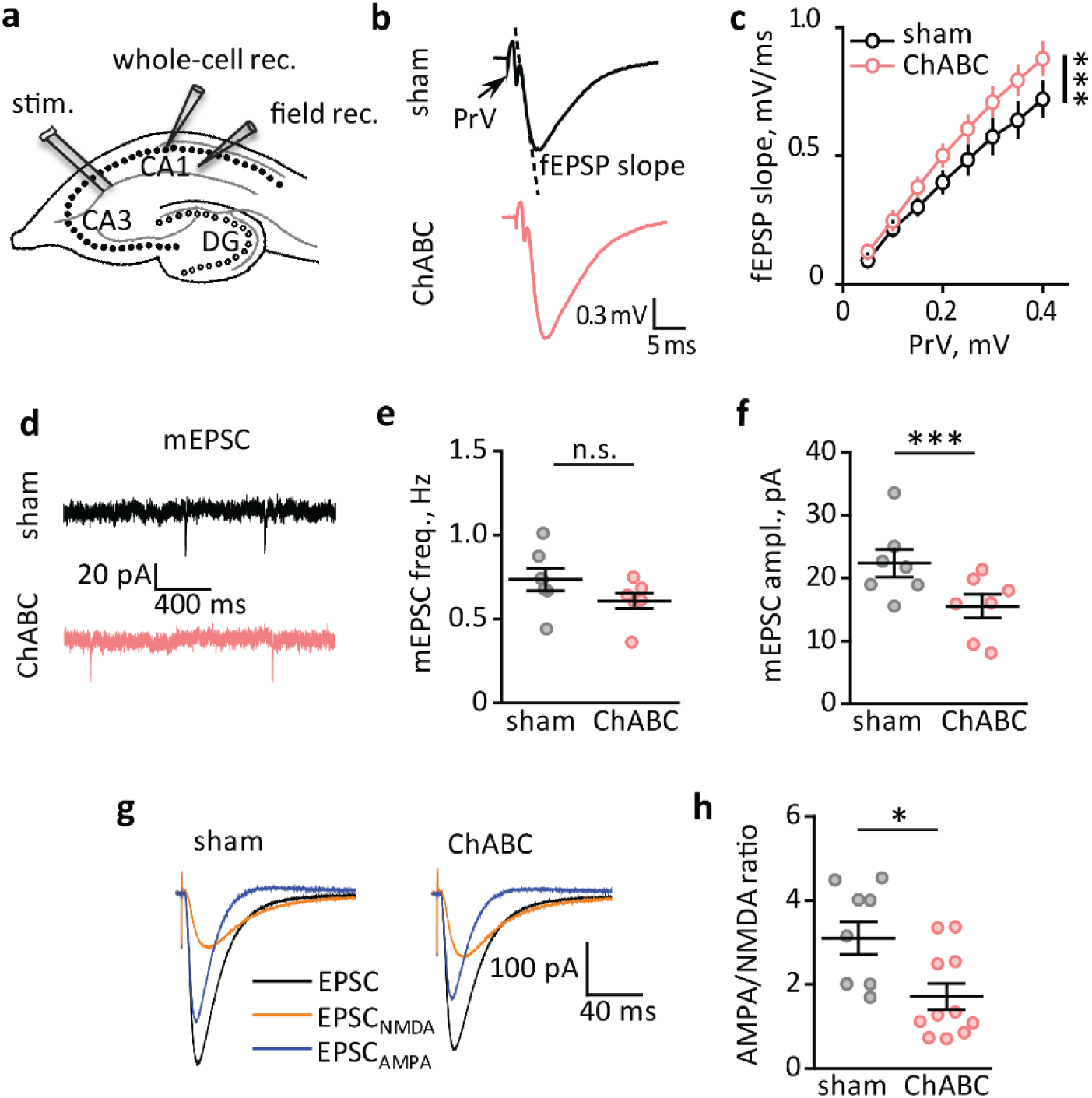
New unpotentiated synapses are formed by existing axons. **a.** Position of the recording and stimulating electrodes in the CA1 region of hippocampal slices. A stimulating electrode (stim.) was placed in the *stratum radiatum* at the border of CA1 to activate Schaffer collaterals. Whole-cell recordings were performed with one stimulating electrode (whole-cell rec.) from CA1 pyramidal neurons. Field potential recordings (PrV and fEPSP) were obtained with an extracellular glass electrode (field rec.) placed at the CA1 *stratum radiatum*. **b.** Representative traces of field potentials (PrV and fEPSP). Arrow points to PrV. The dashed line indicates the fEPSP slope. **c.** The averaged input-output curves or relationships of fEPSP slope to PrV amplitude. **d.** Representative traces of mEPSCs. **e,f.** Summary of the mIPSC frequency (*e*) and mean amplitude (*f*). **g.** Evoked EPSCs were recorded in Mg^2+^ free extracellular solution. At baseline conditions, EPSCs consisted of AMPA and NMDA receptor mediated components (EPSC, *black trace*). Application of NBQX blocked AMPA receptors and revealed NMDA receptor mediated EPSCs (EPSC_NMDA_, *orange trace*). Then, EPSC_NMDA_ was subtracted from EPSC to obtain the AMPA receptor mediated EPSC (EPSC_AMPA_, *blue trace*). **h.** The ratio of the EPSC_AMPA_ to EPSC_NMDA_ amplitude. *Black traces and circles –* recordings in sham-treated slices; *pink traces and circles* - recordings in ChABC-treated slices. The data are presented as the mean ± SEM. n.s. *p* > 0.05; \**p* < 0.05; \*\*\**p* < 0.001; two-way RM ANOVA (panel *c*); two-tailed two-sample *t*-test (panels *e, f,h*).

### New synapses are unpotentiated

The decrease seen in the mean mEPSC amplitude is consistent with the observation that unpotentiated synapses have lower quantal amplitude than potentiated ones (Kullmann and Nicoll, 1992). If the number of unpotentiated synapses increases following ChABC treatment, the ratio of AMPA receptor-mediated to NMDA receptor-mediated currents (AMPA/NMDA ratio) should decrease, reflecting the lower number of AMPA receptors in unpotentiated synapses (Kauer et al., 1988; Lu et al., 2001). Indeed, we observed a significantly smaller AMPA/NMDA ratio in ChABC treated slices (sham: 3.1 ± 0.4, *n* = 9; ChABC: 1.7 ± 0.3, *n* = 11; *p* = 0.012, two-sample *t*-test; Fig. 2g,h).

The mechanism of spontaneous and action potential-dependent vesicular release differ (Kaeser and Regehr, 2014). To determine the mode of release that is affected by ChABC treatment, we analyzed the paired-pulse ratio (PPR) of evoked EPSCs at different inter-stimulus intervals (ISI, Fig. S2a,b). ChABC did not have a significant effect on PPR (*F*(1,40) = 0.36, *p* = 0.6, two-way RM ANOVA; Table S1, Fig. S2b), suggesting that action potential-dependent glutamate release is not affected by ECM attenuation. In addition, we characterized the lateral mobility of postsynaptic AMPA receptors in the membranes of spines. It has been reported that hyaluronidase-mediated attenuation of ECM accelerates the lateral diffusion of postsynaptic AMPA receptors in cultured neurons (Frischknecht et al., 2009). Accelerated movement of AMPA receptors allows for quick replacement of receptors that have been desensitized by synaptic glutamate uncaging and therefore expediates the recovery of the AMPA receptor-mediated current. However, the paired-pulse depression of the glutamate uncaging-induced AMPA receptor currents did not change significantly after ChABC treatment (sham: 0.91 ± 0.04, *n* = 7; ChABC: 0.92 ± 0.03, *n* = 7X; *p* = 0.99, two-sample *t*-test, Fig. S2c-e). This suggests that ECM attenuation in hippocampal slices does not affect the lateral diffusion of AMPA receptors.

### ECM attenuation suppresses LTP

The appearance of additional unpotentiated synapses after ChABC treatment should increase the magnitude of LTP. To test this, we recorded EPSPs in CA1 pyramidal neurons and induced LTP by 5 episodes of theta-burst stimulation of Schaffer collaterals (TBS, Fig. 3a). Surprisingly, LTP was not enhanced but completely suppressed after ChABC treatment (sham: 181 ± 22 % of baseline,n = 8; ChABC: 109 ± 12 % of baseline, *n* = 6; *p* = 0.013, two-sample *t*-test; Fig. 3b,c). To understand the mechanism of such suppression, we analyzed cellular responses to individual, ten-burst episodes of theta-burst stimulation (Fig. 3d). In sham-treated slices, we observed a progressive increase in the number of action potentials from the first to the fifth episode. In the first episode, the number of action potentials in ChABC-treated slices was indistinguishable from that in the sham; however, from the second episode onward, the number of action potentials in the ChABC-treated slices was reduced when compared to sham slices (*F*(1,500) = 183.58, *p* < 0.001, two-way RM ANOVA; Table S1, Fig. 3e). Similar changes were observed in the slope of burst EPSPs. The slope progressively increased during subsequent theta-bursts in the sham-treated slices, but this progressive increase was not apparent after ChABC treatment (*F*(1,500) = 278.44, *p* < 0.001, two-way RM ANOVA; Table S1, Fig. 3f).

**Figure 3.**
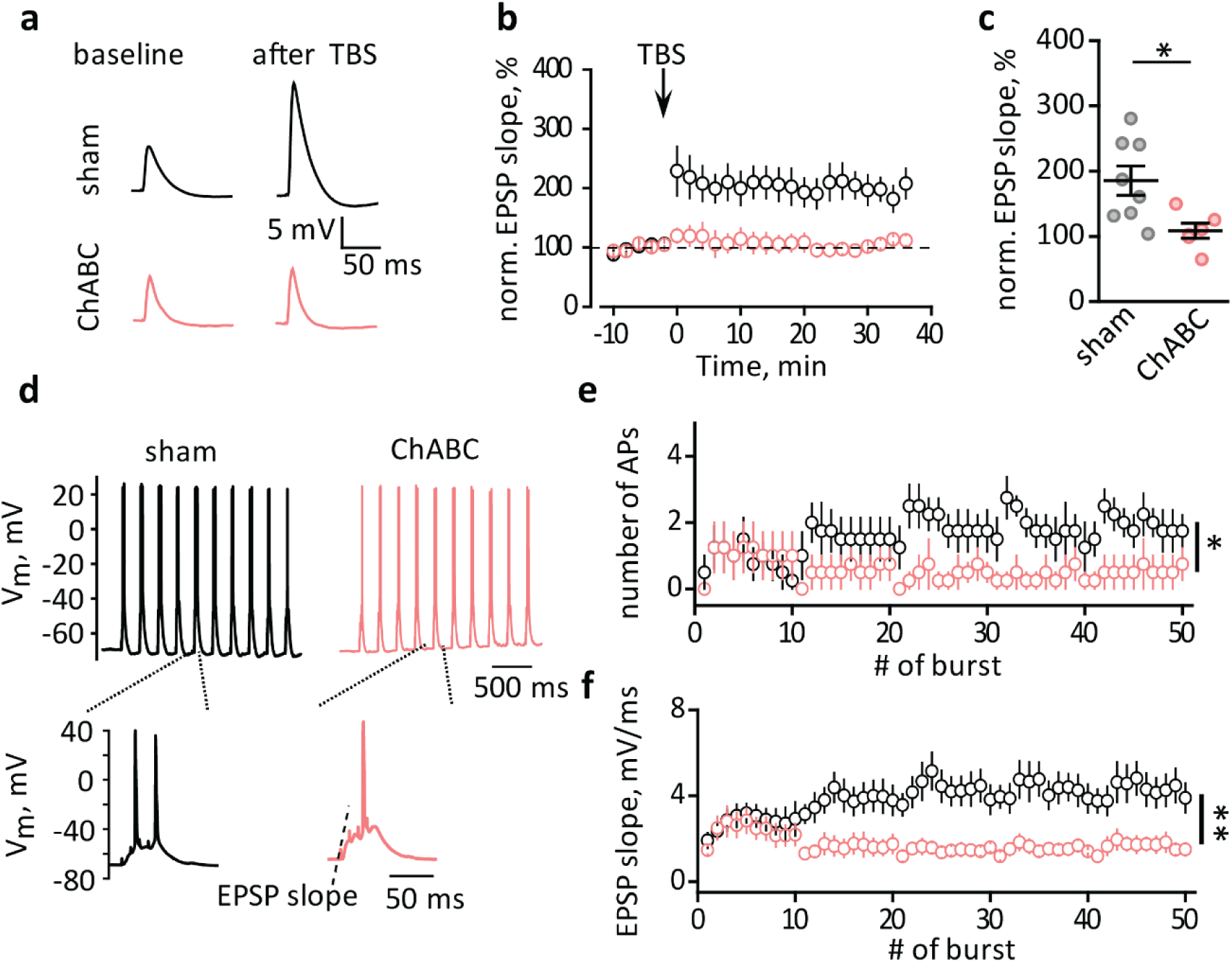
ECM attenuation suppresses LTP. **a.** EPSPs recorded in CA1 pyramidal neurons before (baseline) and after TBS. **b.** The time-course of normalized EPSP slopes before and after TBS (arrow, zero time point). **c.** LTP magnitudes were averaged over the last 10 min of recordings presented in *b*. **d.** Cell responses to TBS (first 10 bursts out of 5 repetitions). Insets show the response to a single burst. The dashed line indicates where the EPSP slope was measured. **e,f.** The number of action potentials (APs, *e*) and EPSP slope (*f*) recorded in response to individual bursts during TBS used to induce LTP (5 series by 10 bursts). *Black traces and circles –* recordings in sham-treated slices; *pink traces and circles* - recordings in ChABC-treated slices. The data are presented as the mean ± SEM. n.s. *p* > 0.05; \**p* < 0.05; \*\**p* < 0.01; two-tailed two-sample *t*-test (panel *c*); two-way RM ANOVA (panels *e,f*).

LTP can be influenced by GABAergic inhibition (Chapman et al., 1998; Grover and Yan, 1999), which may increase upon ECM attenuation (Hirono et al., 2018). ChABC treatment did not produce significant changes in the frequency or amplitude of spontaneous and miniature IPCSs (Fig. S3a-f). Likewise, no significant effect was observed on the PPR of evoked IPSCs (Fig. S3g,h). Thus, ECM attenuation did not affect the basal properties of GABAergic synaptic transmission. However, GABAergic signaling may respond differently to theta-burst stimulation. Theta-burst stimulation did not result in the potentiation of IPSCs in CA1 pyramidal neurons (Fig. S4a-c). Lastly, we monitored changes to IPSCs during individual episodes of theta-burst stimulation (Fig. S4d). No significant changes were observed in the normalized IPSP amplitudes after ChABC treatment (Fig. S4e).

### ECM attenuation reduces the excitability of CA1 pyramidal neurons through upregulation of SK channels

The suppression of LTP may indicate that ChABC treatment reduces the excitability of postsynaptic neurons. Indeed, ChABC treatment significantly reduced the number of action potentials evoked in CA1 pyramidal neurons following 500 ms current injections. (*F*(1,100) = 21.99, *p* < 0.001, two-way RM ANOVA; Table S1, Fig. 4a,b). This effect of ECM attenuation remained even after the blockade of glutamatergic and GABAergic ionotropic receptors with 50 µM APV, 50 µM NBQX, and 100 µM picrotoxin (Fig. S5). No significant change in the input resistance (Rin) of cells was observed after ChABC treatment (sham: Rin = 172 ± 20 MΩ, n=6, ChABC: Rin = 162 ± 17 MΩ, *n* = 6; *p* = 0.82, two-sample *t*-test; Fig. 4c and S5c,f). Therefore, tonic conductances mediated by extrasynaptic receptors or plasma membrane channels are not involved in reducing the excitability of pyramidal neurons following ChABC treatment. The observed decrease in cell excitability could be associated with the downregulation of voltage-gated Na^+^ channels or the upregulation of voltage-gated K^+^ channels which would then affect the waveform and threshold of action potentials. However, neither the action potential threshold (sham: −44.5 ± 1.5 mV, *n* = 6; ChABC: −46.2 ± 0.8 mV, *n* = 6; *p* = 0.94, two-sample *t*-test) nor the action potential half-width (sham: 1.49 ± 0.18 ms, *n* = 6; ChABC: 1.55 ± 0.15 ms, *n* = 6; *p* = 0.81, two-sample *t*-test) were affected by ChABC treatment (Fig. S6). An after hyperpolarization (AHP) of increased magnitude is another possible mechanism behind the reduced excitability. Indeed, ChABC treatment significantly increased the AHP recorded in response to an injection of depolarizing current (*F*(1,100) = 25.96, *p* < 0.001, two-way RM ANOVA; Table S1, Fig. 4d).

**Figure 4.**
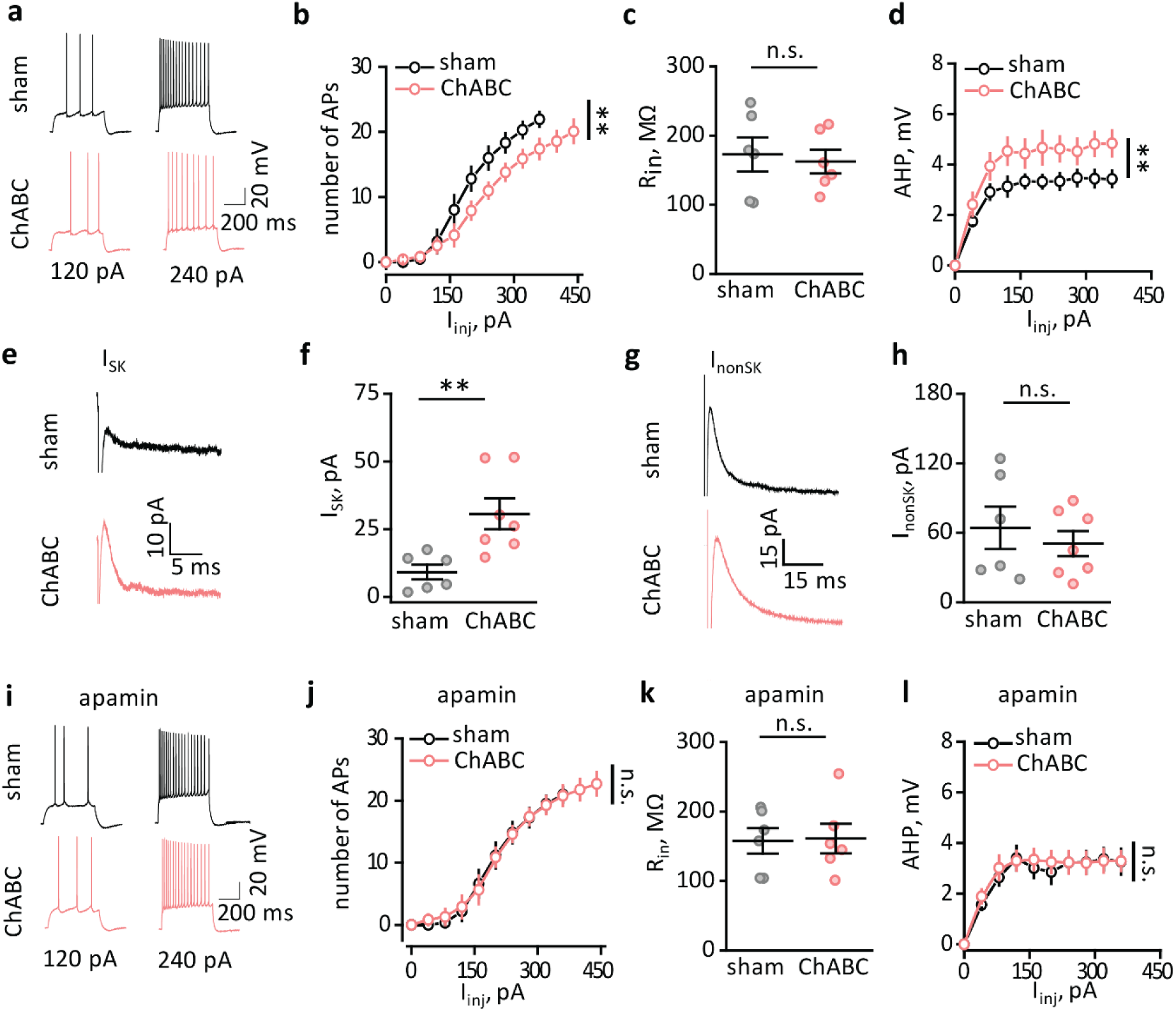
ECM attenuation reduces the excitability of CA1 pyramidal neurons through upregulation of SK channels. **a.** Action potentials elicited by depolarizing current steps (120 pA and 240 pA). **b.** Summary of the number of action potentials (APs) that occur in response to depolarizing steps. **c.** Summary of the input resistance (Rin). **d.** Summary of AHP amplitudes. **e.** Representative traces of I_SK_. **f.** Summary of I_SK_ amplitudes. **g.** Representative traces of I_nonSK_. **h.** Summary of I_nonSK_ amplitudes. **i.** Action potentials (APs) elicited by depolarizing current steps (120 pA and 240 pA) in the presence of the SK channel blocker, apamin. **j.** Summary of the number of APs that occur in response to depolarizing steps in the presence of apamin. **k.** Summary of Rin in the presence of apamin. **l.** Summary of AHP amplitude in the presence of apamin. *Black traces and circles –* recordings in sham-treated slices; *pink traces and circles* - recordings in ChABC-treated slices. The data are presented as the mean ± SEM. n.s. *p* > 0.05; \*\**p* < 0.01; two-way RM ANOVA (panels *b,d,j,l*); two-tailed two-sample *t*-test (panels *c,f,h,k*).

Several types of Ca^2+^-activated K^+^ channels contribute to the AHP (Savic et al., 2001). First, we measured the current mediated by small conductance Ca^2+^-activated K^+^ (SK) channels (I_SK_). We recorded an outward current following a depolarizing step (50 ms, 70 mV) in the presence of 1 µM tetrodotoxin, a Na^+^ channel blocker. Then we applied 100 nM apamin, an SK channel blocker, and thus obtained the remaining, non-SK channel-mediated outward current (I_nonSK_). We obtained I_SK_ by subtracting I_nonSK_ from the total current. ChABC treatment significantly increased I_SK_ (sham: 9.2 ± 2.7 pA, *n* = 6; ChABC: 30.7 ± 5.7 pA, *n* = 7; *p* = 0.003, two-sample *t*-test; Fig. 4e,f) but had no significant effect on I_nonSK_ (sham: 64.3 ± 18.3 pA, *n* = 6; ChABC: 50.7 ± 10.8 pA, *n* = 7; *p* = 0.36, two-sample *t*-test; Fig. 4g,h). Thus, ECM attenuation upregulates SK channels but does not upregulate other Ca^2+^-activated K^+^ channels. This rules out the possible involvement of voltage-dependent Ca^2+^ channels (Kochlamazashvili et al., 2010), as it would affect all Ca^2+^-activated K^+^ channels.

Next, we tested if the upregulation of SK channels is responsible for the reduced excitability of CA1 pyramidal neurons following ChABC treatment. Apamin abolished both the decrease in cell excitability (cell firing in apamin: *F*(1,100) = 0.01, *p* = 0.9, two-way RM ANOVA; Table S1, Fig. 4i,j) and AHP amplification (AHP in apamin: *F*(1,108) = 0.41, *p* = 0.5, two-way RM ANOVA; Table S1, Fig. 4k,l) induced by ChABC treatment. Consistent with the lack of contribution of other Ca^2+^-activated K^+^ channels, the decrease in cell excitability induced by ChABC treatment was preserved in the presence of 5 µM paxilline, a blocker of large conductance Ca^2+^-activated K^+^ (BK) channels (cell firing in paxilline: *F*(8,72) = 13.40, *p* < 0.001, two-way RM ANOVA; Table S1,Fig. S7)

### Blockade of SK channels reveals enhanced LTP upon ECM attenuation

Next, we tested if a blockade of SK channels could also rescue the suppression of LTP after ChABC treatment. In agreement with the increased proportion of unpotentiated synapses, LTP was not only rescued, but significantly enhanced in the presence of apamin (sham + apamin: 155 ± 9 % of baseline, *n* = 8; ChABC + apamin: 266 ± 31 % of baseline, *n* = 7; *p* = 0.003, two-sample *t*-test; Fig. 5a-c). Notably, the number of action potentials and the increased EPSP slope during theta burst stimulation were rescued, but not enhanced, by apamin (number of action potentials: *F*(1,600) = 3.17, *p* = 0.08, two-way RM ANOVA; increase in EPSP slope: *F*(1,600) = 0.97, *p* = 0.33, two-way RM ANOVA; Table S1, Fig. 5e,d).

**Figure 5.**
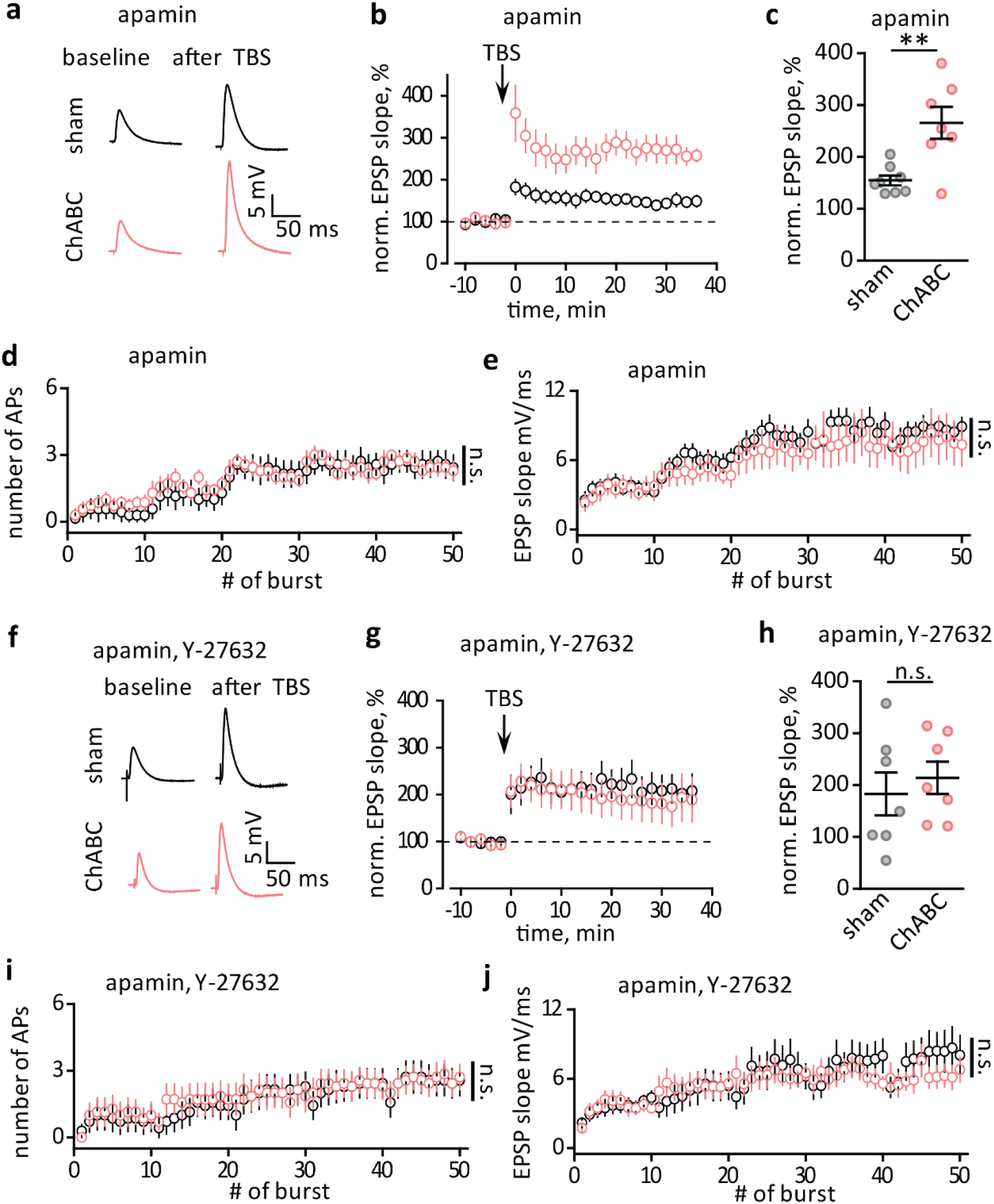
Blockade of SK channels reveals ROCK pathway-dependent enhancement of LTP. **a.** EPSPs recorded in CA1 pyramidal neurons before (baseline) and after TBS in the presence of apamin. **b.** The time-course of normalized EPSP slopes before and after TBS (arrow, zero time point) in the presence of apamin. **c.** LTP magnitudes averaged over the last 10 min of recordings presented in *b*. **d,e.** The number of action potentials (APs, *d*) and the EPSP slope (*e*) recorded in response to individual bursts during TBS used to induce LTP (5 series by 10 bursts) in the presence of apamin. **f.** EPSPs recorded in CA1 pyramidal neurons before (baseline) and after TBS in the presence of apamin and the ROCK inhibitor Y-27632. **g.** The time-course of normalized EPSP slopes before and after TBS (arrow, zero time point) in the presence of apamin and Y-27632. **h.** LTP magnitudes averaged over the last 10 min of recordings presented in *b*. **i,j.** The number of APs (*i*) and the EPSP slope (*j*) recorded in response to individual bursts during TBS used to induce LTP (5 series by 10 bursts) in the presence of apamin and Y-27632. *Black traces and circles –* recordings in sham-treated slices; *pink traces and circles* - recordings in ChABC-treated slices. The data are presented as the mean ± SEM. n.s. *p* > 0.05; \*\**p* < 0.01; two-tailed two-sample *t*-test (panels *c,h*); two-way RM ANOVA (panels *d,e,i,j*).

If the enhanced LTP can be attributed to the enlargement of thin spines, it should be associated with pathways involved in cytoskeletal remodeling. These pathways require activation of small GTPases, such as RhoA (Murakoshi et al., 2011). RhoA controls spine remodeling through Rho-associated protein kinase (ROCK) (Nakayama et al., 2000). Indeed, the ROCK inhibitor Y-27632 (10 µM) abolished the enhancement of LTP observed in ChABC treated slices in the presence of apamin (sham + apamin + Y-27632: 182.8 ± 41.4 % of baseline, *n* = 7; ChABC + apamin + Y-27632: 213.8 ± 31.0 % of baseline, *n* = 7; *p* = 0.56, two-sample *t*-test; Fig. 5f-h). The number of action potentials and the increased EPSP slope during theta burst stimulation were not affected by the ROCK inhibitor (number of action potentials: *F*(1,600) = 1.81, *p* = 0.18; increase in EPSP slope: *F*(1,600) = 2.93, *p* = 0.1, two-way RM ANOVA; Table S1, Fig. 5i,j). Thus, the observed LTP consisted of two components: ROCK-dependent and ROCK-independent LTP. The ROCK-dependent component is linked to spine enlargement and is promoted by ECM attenuation.

## Discussion

The ECM holds neighboring cells together and stabilizes synapses (Dansie and Ethell, 2011; Levy et al., 2014). However, the ECM also shields the cell surface making it more difficult for new synapses to form (Bikbaev et al., 2015; Orlando et al., 2012). The effect of ECM removal on dendritic spines has been previously shown in neuronal culture and organotypic slices, where it increases spine motility, spine extension, and network rewiring (Bikbaev et al., 2015). Physiologically, the ECM is quite diffuse at early developmental stages, allowing for neuronal network wiring and rewiring (Frischknecht and Gundelfinger, 2012). In the adult brain, the ECM stabilizes existing dendritic spines and restricts the appearance of new ones. Hence, the morphological plasticity of dendritic spines and synaptic rewiring requires activity-dependent recruitment of multiple extracellular proteases, such as ADAMTS4/5/15, MMP9, and neurotrypsin, that produce localized degradation of ECM (Bijata et al., 2017; Ferrer-Ferrer and Dityatev, 2018). Injection of ChABC has been demonstrated to promote axonal sprouting in the pericontusion cortical region *in vivo* (Harris et al., 2010). In this study, we observed that the ECM attenuation induced by ChABC triggers the formation of new synapses between existing axons and new thin dendritic spines in the hippocampus.

The thin dendritic spines that appear upon ECM attenuation are ‘learning spines.’ The name indicates the ability of these spines to undergo further morphological maturation during synaptic enhancement and conversion to mushroom or ‘memory’ spines (Bourne and Harris, 2007). Mushroom spines host more AMPA receptors, which makes their synapses stronger (Matsuzaki et al., 2001; Nimchinsky et al., 2004). Smaller spines with thinner necks exhibit a larger increase in Ca^2+^ concentration during LTP induction; hence, they are more likely to be potentiated (Noguchi et al., 2005). During LTP, new AMPA receptors get inserted into the postsynaptic density (PSD) (Bredt and Nicoll, 2003; Malinow and Malenka, 2002). Both the spine and PSD increase in size, leading to spine ‘maturation’ (Bosch et al., 2014).

Our morphological observations did not reveal any significant changes in the density of mushroom spines after ChABC treatment. Electrophysiologically, we found that both the mean amplitude of mEPSC and the AMPA/NMDA ratio decreased. Thus, ECM attenuation increased the proportion of unpotentiated thin spines. This would be expected to enhance LTP, as the magnitude of LTP depends on the ratio of unpotentiated to potentiated synapses (Govindarajan et al., 2011; McNaughton and Morris, 1987); surprisingly, LTP was suppressed in ChABC treated slices. This LTP suppression could not be attributed to changes in the efficiency of glutamatergic or GABAergic synaptic transmission following ECM attenuation. mEPSC frequency; the paired-pulse ratio of evoked EPSCs; sIPSC/mIPSC frequency and amplitude; and the PPR of evoked IPSCs appeared to be unchanged after ChABC treatment. It has been reported that ECM limits lateral diffusion of AMPA receptors along the plasma membrane and hence regulates synaptic strength in neuronal cultures (Favuzzi et al., 2017; Frischknecht et al., 2009). However, we did not find evidence that attenuation of ECM with ChABC affects the lateral diffusion of AMPA receptors in hippocampal slices. This finding is consistent with another report which suggested that the ECM does not regulate the lateral diffusion of AMPA receptors in aspiny neurons (Klueva et al., 2014).

We hypothesized that increased membrane conductance and decreased postsynaptic neuron excitability may be responsible for the observed suppression of LTP. The ECM protein vitronectin regulates inactivating potassium currents (*IA*) in embryonic mouse hippocampal neurons (Vasilyev and Barish, 2003); brevican controls the localization of potassium channels in parvalbumin-positive interneurons (Favuzzi et al., 2017); and hyaluronic acid regulates hippocampal synaptic plasticity by modulating postsynaptic L-type Ca^2+^ channels (Kochlamazashvili et al., 2010). In this study, ChABC treatment decreased the excitability of CA1 pyramidal neurons. This decrease was not accompanied by any change in action potential threshold or waveform, ruling out an effect of ECM attenuation on Na^+^ or K^+^ channels. The input resistance of cells was likewise unaffected, indicating a lack of effect on baseline membrane conductances. The reduced excitability of CA1 pyramidal neurons was mediated by the upregulation of SK channels. Indeed, SK channels are known to cause early spike frequency adaptation and contribute to AHP in CA1 pyramidal neurons (Chen et al., 2014). SK channels have also been reported to curtail Ca^2+^ influx in dendritic spines and shafts during neuronal firing and to reduce synaptic plasticity (Griffith et al., 2016; Jones and Stuart, 2013; Jones et al., 2017). The SK channel blocker apamin increases LTP and facilitates learning (Adelman et al., 2012). Here, an application of apamin restored both the excitability of cells and LTP that were suppressed by ECM attenuation.

It is unknown if the upregulation of SK channels can be mediated directly by ECM attenuation. Currently, there is no evidence that components of the ECM interact with SK channels. However, SK channels are directly coupled to NMDA receptors in dendritic spines (Luján et al., 2009). Thus, the appearance of new dendritic spines may potentially increase the total number of SK channels in the neuron. In other words, each dendritic spine adds a ‘quantum’ of SK channels, progressively decreasing the excitability of the cell. In fact, during cerebellar development, the formation of new synapses is associated with a progressive increase in the expression of SK channels in Purkinje cells (Ballesteros-Merino et al., 2014). It is possible that the upregulation of SK channels alongside an increase in the number of excitatory synapses serves as a protective mechanism against overexcitation of the neuronal network, which can lead to excitotoxicity and epilepsy. A similar mechanism has been revealed during LTP induction in CA1 pyramidal neurons, where LTP triggers upregulation of hyperpolarization-activated cation (*h*) channels, thereby decreasing overall cell excitability and balancing the additional excitation stemming from increased synaptic strength (Fan et al., 2005; Wu et al., 2012). Thus, in order to maintain network stability, CA1 pyramidal neurons possess mechanisms to balance the excitability associated with increased synaptic strength and/or an increased number of excitatory synapses.

During LTP induction, thin dendritic spines undergo morphological changes that require remodeling of the cytoskeleton and actin polymerization (Fukazawa et al., 2003; Krucker et al., 2000; Okamoto et al., 2004). First, Ca^2+^ enters through NMDARs and voltage-gated Ca^2+^ channels and then activates Ca^2+^/calmodulin kinase II (CaMK II) (Bosch et al., 2014; Furuyashiki et al., 2002). Next, CaMK II activates the RhoA-ROCK pathway, which controls the enlargement of spines (Hedrick et al., 2016; Murakoshi et al., 2011; Rex et al., 2009). The upregulation of SK channels reduces Ca^2+^ influx during LTP induction and prevents activation of this pathway (Griffith et al., 2016). When SK channels were blocked with apamin, the full strength of LTP was unleashed. Consistent with a higher proportion of unpotentiated synapses, the magnitude of LTP was increased by ECM attenuation. Blocking ROCK abolished the enhancement of LTP in ChABC treated slices, suggesting that the mechanism behind this enhancement is related to the potentiation of synapses on thin spines. However, the ROCK blockade did not completely abolish LTP. This finding suggests that a certain amount of LTP is independent of the RhoA-ROCK pathway and that this form of LTP is not affected by ECM attenuation.

In summary, we conclude that ECM suppresses the growth of new spines in hippocampal pyramidal neurons (Fig.6). When the ECM is attenuated by exogenous treatment or by endogenous metalloproteinases new synaptic contacts are formed on thin dendritic spines. The appearance of new spines parallels the upregulation of SK channels that leads to reduced excitability of neurons and suppresses LTP. This phenomenon may represent a mechanism of neuronal network stabilization but may also lead to impaired learning and memory.

**Figure 6.**
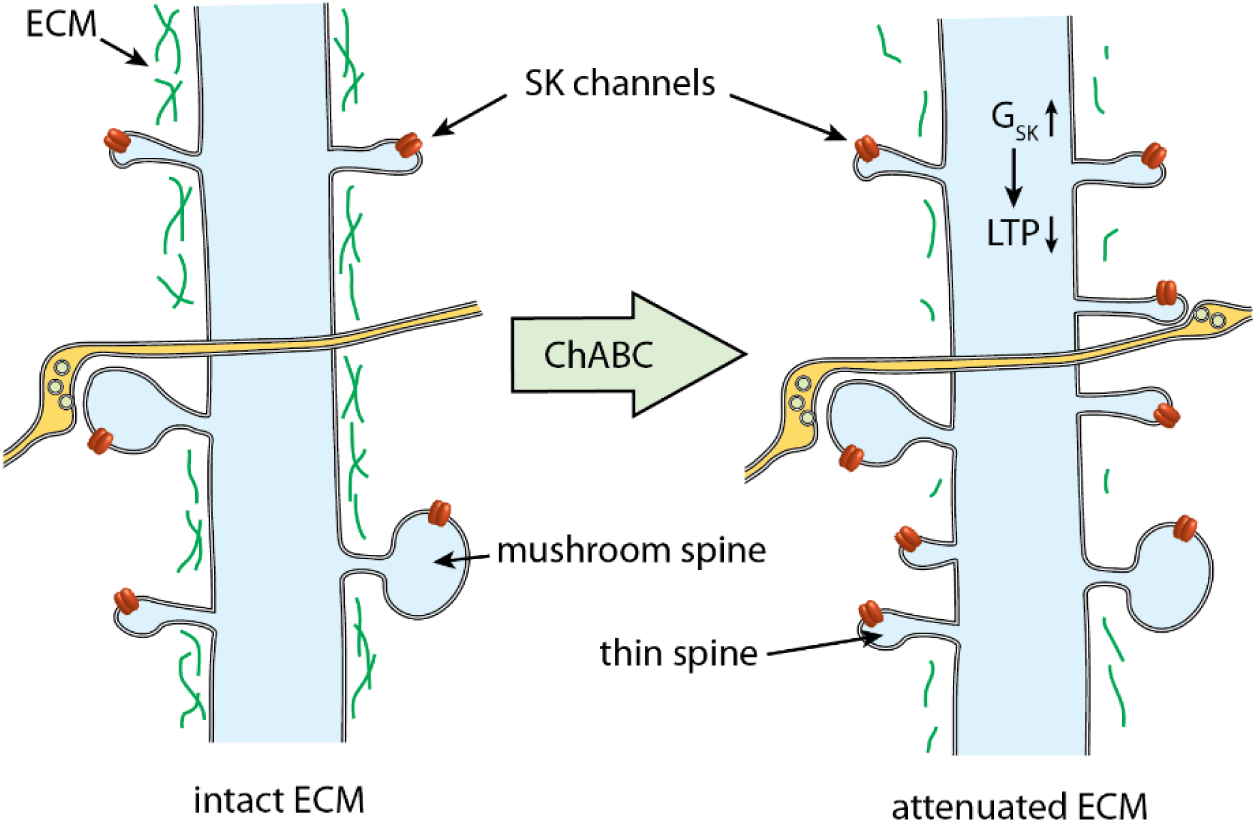
A schematic showing the effect of ECM attenuation on a dendrite of a CA1 pyramidal neuron. The ECM forms a meshwork (*green*) around the dendrite (*blue*), preventing the formation of new spines. When the ECM is attenuated with ChABC, new thin spines emerge. These spines form synaptic contacts with existing axons, Schaffer collaterals (*yellow*). The number of mature mushroom spines does not change. The appearance of new spines adds extra SK channels. An increase in the overall SK channel conductance (G_SK_) decreases neuronal excitability and suppresses LTP.

## Methods

### Animals

All experiments were performed in 4- to 6-week-old C57BL/6J mice. Animal procedures were conducted in accordance with regulations and ethical animal research standards defined by the ethical committee of the Lobachevsky State University of Nizhny Novgorod.

### Hippocampal slice preparation

Animals were killed by cervical dislocation and then decapitated. The brains were exposed and chilled with ice-cold solution containing (in mM) 87 NaCl, 2.5 KCl, 7 MgCl2, 1.25 NaH2PO4, 26.2 NaHCO3, 0.5 CaCl2, 25 D-glucose, and 50 sucrose. Hippocampi from both hemispheres were isolated, and 350 μm-thick transverse slices were cut with a vibrating microtome (Microm HM 650V, Thermo Fisher Scientific, or VT1200S, Leica). Slices were incubated in a 3-ml chamber for 2 hours at 37°C in a solution containing (in mM) 113 NaCl, 2.38 KCl, 1.24 MgSO4, 0.95 NaH2PO4, 24.9 NaHCO3, 1 CaCl2, 1.6 MgCl2, 27.8 D-glucose and 0.2% bovine serum albumin (sham) or in the same solution supplemented with 0.2 U/ml of protease-free chondroitinase ABC (ChABC) from *Proteus vulgaris* (Amsbio, UK). Next, the slices were transferred to a recording chamber and were continuously perfused with a solution containing (in mM) 119 NaCl, 2.5 KCl, 1.3 MgSO4, 1 NaH2PO4, 26.2 NaHCO3, 2.5 CaCl2, and 11 D-glucose. All solutions were saturated with 95% O2 and 5% CO2. The osmolarity was 295 ± 5 mOsm.

### Electron microscopy

Acute slices of two animals were treated with either sham or ChABC solutions and were placed into a solution containing 3% paraformaldehyde and 0.5% glutaraldehyde in 0.1 M Na-cacodylate buffer (pH 7.2–7.4) at room temperature. The slices were further fixed for electron microscopy by immersion in 2.5% glutaraldehyde in the same 0.1 M Na-cacodylate buffer for 24 h. The tissue was postfixed with 1% osmium tetroxide and 0.01% potassium dichromate in the same buffer for 1–2 h at room temperature. The tissue was then dehydrated in graded aqueous solutions of ethanol from 40% to 96% (each for 10 min) and then 100% acetone (three changes, each for 10 min). Specimens were infiltrated with a mixture of 50% epoxy resin and 50% pure acetone for 30 min at room temperature. Each slice was placed on an Aclar film and covered with a capsule containing pure epoxy resin (Epon 812/AralditeM epoxy resins) at 60°C for 1 h and polymerized overnight at 80°C. The epoxy resin blocks with tissue slices were coded and all further analyses were carried out by an investigator blinded to the experimental status of the tissue. The embedded slices on the block surface were trimmed with a glass knife along the entire surface of the hippocampal slice, and 1-µm thick sections were cut. A trapezoidal area was prepared with a glass knife with one side of 250–350 µm in length, and that included the CA1 hippocampal area. Images of serial sections (60 nm) were cut with a Diatome diamond knife and allowed to form a ribbon on the surface of a water/ethanol solution (2%–5% ethanol in water) in the knife bath and collected using Pioloform-coated slot copper grids. Sections were counterstained with saturated ethanolic uranyl acetate, followed by lead citrate, and then placed in a rotating grid holder to allow for the uniform orientation of sections on adjacent grids in the electron microscope. Serial sections in the middle of the *stratum radiatum* at a location ~125-150 µm from the CA1 pyramidal cell body layer were obtained at a magnification of 4000x in a JEOL 1010 electron microscope. Each image was around 20.8 by 11.2 µm^2^. Up to 150 serial sections per series were photographed for reconstruction.

### Stereological and 3D reconstruction analysis

Images in the series were aligned automatically using homemade software. Further analysis was done in Reconstruct software (http://synapseweb.clm.utexas.edu/software-0). The unbiased brick method (Fiala and Harris, 2001) was used for correct sampling from the total synapse population being presented in the acquired volume of CA1 *stratum radiatum*. Each image stack was divided into 5 parts, 30 sections in depth. Unbiased bricks 2 µm x 2 µm in size and 10 sections (0.6 µm) in depth were positioned randomly into each part of the image stack using a sampling grid of X: 8 µm, Y: 6 µm in size. In total, 30 unbiased bricks were analyzed (Fig.S1). Three sides of the brick were inclusion planes, and the other three sides were exclusion planes. Postsynaptic densities (PSD) were used as markers for dendritic spines. Those PSDs that were completely inside the brick or touching only the inclusion planes were counted, whereas PSDs that made any contact with the exclusion planes were excluded. All countable dendritic spines, their PSDs, and presynaptic boutons were traced and generated meshed 3D objects, of which the volume and surface area figures were used for further morphological analysis. Approximately 120-180 spines/PSDs were reconstructed per animal. 3D reconstructions of the selected dendritic spines were imported to 3D Studio Max 2016 software for rendering and subsequent rotation to display the optimal view of the reconstructed structures.

### Electrophysiology

Presynaptic fiber volley (PrV) and field EPSP (fEPSP) were simultaneously recorded with a glass electrode (3 - 5 MΩ resistance) filled with the extracellular solution and placed in CA1 *stratum radiatum* (field rec. at Fig. 2a). Stimulation of Schaffer collaterals was performed with a bipolar stainless-steel electrode (FHC, Bowdoinham, ME, USA) at a distance of more than 150 μm from the recording site (stim. at Fig. 2a). The amplitude of the baseline stimulation frequency was 0.05 Hz, which induced a half-maximal response. The stimulus intensity was varied to obtain the relationship between the fEPSP slope and PrV amplitude (the input-output curve). PrV amplitudes were binned with 0.05 mV bin size in the range from 0 to 0.45 mV and the mean fEPSP slope was calculated for each bin.

Whole-cell recordings from CA1 pyramidal neurons were obtained with glass electrodes (3–5 MΩ resistance). mEPSCs were recorded using an intracellular solution containing (in mM): 130 KCH3SO3, 8 NaCl, 10 Na phosphocreatine, 10 HEPES, 2 EGTA, 3 Na L-ascorbic acid, 10 HEPES, 0.4 NaGTP, 2 MgATP, and 5 QX314 Br (pH adjusted to 7.2 with KOH and osmolarity adjusted to 290 mOsm.). The membrane potential was clamped at −70 mV. The extracellular solution contained μM picrotoxin, 200 μM (S)-a-methyl-4-carboxyphenyglycine (MCPG), 5 μM CGP52432 and 1 μM tetrodotoxin to block GABAA, mGluR, GABAB receptors and action potentials, respectively.

Spontaneous inhibitory postsynaptic currents (sIPSCs) were recorded using pipette solution containing (in mM): 120 CsCl, 8 NaCl, 0.2 MgCl2, 10 HEPES, 2 EGTA, 0.3 NaGTP, 2 MgATP, and 5 Qx314 Br (pH adjusted to 7.2 with CsOH and osmolarity adjusted to 290 mOsm). The membrane potential was clamped at −70 mV. The extracellular solution contained 50 μM APV, 25 μM NBQX, 200 μM MCPG, and 5 μM CGP52432 to block NMDA, AMPA, mGluR, and GABAB receptors, respectively. For mIPSCs recordings, 1 μM tetrodotoxin was added to this drug cocktail. Schaffer collaterals were stimulated at 20, 50, 100, and 150 ms interstimulus intervals to measure the PPR of the evoked IPSCs.

EPSCs mediated by AMPA (EPSC_AMPA_) and NMDA (EPSC_NMDA_) receptors were recorded using an intracellular solution containing (in mM): 130 KCH_3_SO_3_, 8 NaCl, 10 Na phosphocreatine, 10 HEPES, 2 EGTA, 3 Na L-ascorbic acid, 10 HEPES, 0.4 NaGTP, 2 MgATP, and 5 QX314 Br (pH adjusted to 7.2 with KOH and osmolarity adjusted to 290 mOsm) in Mg^2+^ free extracellular solution in the presence of 100 μM picrotoxin, 200 μM MCPG, and 5 μM CGP52432. The membrane potential was clamped at −70 mV. In this set of experiments, CA3-CA1 connections were cut to prevent over-excitation. Shaffer collaterals were stimulated using extracellular bipolar microelectrodes. The stimulation strength was adjusted to evoke composite EPSCs with amplitudes of ~300 pA, and 15 evoked EPSCs were averaged. Then, the EPSC_NMDA_ were pharmacologically isolated by applying 25 μM NBQX, an AMPA receptor blocker. The EPSC_AMPA_ was obtained by subtraction of EPSC_NMDA_ from the composite EPSC. The AMPA/NMDA ratio was calculated as a ratio of EPSC_AMPA_ to EPSC_NMDA_ amplitudes. The EPSC paired-pulse ratio was obtained at several inter-pulse intervals under the same conditions but without NBQX and in the presence of Mg^2+^ in the extracellular solution.

Cell excitability was measured using a pipette solution containing (in mM) 140 K-gluconate, 8 NaCl, 0.2 CaCl2, 10 HEPES, 2 EGTA, 0.5 NaGTP, and 2 MgATP (pH adjusted to 7.2 with KOH and osmolarity adjusted to 290 mOsm). Membrane potential was manually adjusted at −70 mV in current-clamp mode, and 500-ms current steps were injected with increasing amplitudes from - 80 pA to 440 pA in steps of 40 pA. The first steps which did not trigger action potentials (−80 pA to +80 pA range) were used to plot IV-curves (not presented) and calculate the cell input resistance using Ohm’s law. The action potential threshold and half-width were obtained at the first current steps at which action potentials appeared (rheobase). The threshold was calculated as the value of the membrane potential corresponding to the speed of change of the membrane potential (ΔV_m_/Δt) of 20 mV/ms. Half-width was calculated as the duration of the action potential at the voltage halfway between the threshold and the action potential peak. The afterhyperpolarization (AHP) was calculated as the amplitude of a negative peak after a 500 ms depolarizing step relative to the baseline before the depolarizing step.

I_SK_ and I_nonSK_ were measured with intracellular solution containing (in mM) 140 K-gluconate, 8 NaCl, 0.2 CaCl_2_, 10 HEPES, 2 EGTA, 0.5 NaGTP, and 2 MgATP (pH adjusted to 7.2 with KOH and osmolarity adjusted to 290 mOsm) in the presence of 1 μM tetrodotoxin. Membrane potential was clamped at −70 mV, and 500 ms depolarizing steps to +40 mV were delivered to induce potassium current. I_SK_ was blocked and I_nonSK_ was pharmacologically isolated with 100 nM apamin. Then, I_nonSK_ was subtracted from the total potassium current to obtain I_SK_.

### Long-term potentiation (LTP)

LTP was recorded in single neurons in current-clamp mode with glass electrodes filled with intracellular solution containing (in mM): 132.5 K gluconate, 8 NaCl, 10 Na phosphocreatine, 10 HEPES, 3 Na L-ascorbic acid, 0.5 NaGTP, and 2 MgATP (pH adjusted to 7.2 with KOH and osmolarity adjusted to 290 mOsm). Membrane potential was manually adjusted to −70 mV. The stimulation strength was adjusted to obtain EPSP amplitudes in the range of 5-7 mV. Theta-burst stimulation (TBS) was applied five times (at 0.2Hz) to induce LTP and consisted of 4 stimuli at 100 Hz delivered 10 times at 5 Hz. The baseline recording lasted 10 min before TBS. LTP was recorded for 40 min after TBS. The mean EPSP slope recorded before TBS was taken as 100%. LTP values were calculated from 30 to 40 min after TBS.

The effect of TBS on evoked IPSCs was recorded with a pipette solution containing (in mM): 130 CsCH_3_SO_3_, 8 NaCl, 10 Na phosphocreatine, 10 HEPES, 2 EGTA, 3 Na L-ascorbic acid, 10 HEPES, 0.4 NaGTP, 2 MgATP, and 5 QX314 Br (pH adjusted to 7.2 with KOH and osmolarity adjusted to 290 mOsm). No glutamate receptor blockers were added to the extracellular solution. The membrane potential was clamped at the reversal potential of glutamatergic currents, 0 mV. TBS was applied five times at 0.2Hz.

### Two-photon imaging and glutamate uncaging

Cells were filled with 50 µM Alexa 594 for at least 10 min prior to recordings to ensure uniform diffusion of the dye. Two-photon imaging was performed with a two-scanner FV1000-MPE laser-scanning microscope (Olympus, Japan) equipped with a mode-locked (<140 fs pulse width) tunable 720–930 nm laser Chameleon XR (Coherent, USA). Alexa 594 was excited at a wavelength of 830 nm and its fluorescence was chromatically separated and detected with a photomultiplier (PMTs). The bright Alexa Fluor 594 emission identified oblique apical dendrites (about 150 mm from the soma) and their spines. 600 μM of 4-methoxy-7-nitroindolinyl-caged-L-glutamate (MNI-caged-L-glutamate;) was added to the bath for glutamate uncaging. Single-photon uncaging was carried out using 5–10 ms laser pulses (405 nm diode laser; FV5-LD405; Olympus, Japan) with ‘‘point scan’’ mode in Fluoview (Olympus). The uncaging spot was located at the edge of the spine head. The strength of uncaging was adjusted to achieve an uncaging-induced synaptic current amplitude of 150-250 pA. AMPA current (IAMPA) was recorded in the presence of 50 μM APV. The desensitization of AMPA receptors was tested as IAMPA PPR with a 100 ms interpulse interval.

### Drugs and chemicals

All drugs were kept as concentrated stock solutions at −20°C. Apamin, paxilline, CGP55845, Y-27632 were added from the beginning of experiments. MNI-caged glutamate was applied 10 min before the recording. Picrotoxin, MCPG, D-APV, NBQX, tetrodotoxin, CGP55845, apamin, paxilline, and MNI-caged glutamate were purchased either from Tocris Cookson (Bristol, UK) or Sigma Aldrich (Darmstadt, Germany). Y-27632 (10 μM) was added to sham- and ChABC-treated slices during the whole incubation period. Alexa Fluor 594 was obtained from Invitrogen (Carlsbad, CA, USA).

### Data Analysis

Electrophysiological data were analyzed using WinWCP, Mini analysis (6.0.2, Synaptosoft, USA), and Clampfit (9.0 Axon Instruments Inc.; Union City, CA, USA). Statistical analysis was performed with Excel (Microsoft, USA), Origin 8 (OriginLab Corp.), SigmaPlot (12.3 Systat Software Inc., USA), and Matlab 2012b (The MathWorks Inc., USA). All data are presented as the mean ± SEM, and differences between groups were tested with unpaired *t*-test and two-way RM ANOVA, as stated in the text. p > 0.05 was considered significant.

## Acknowledgments

The authors are grateful to Prof. Alexander Dityatev for valuable comments, Dr Inseon Song for help with electrophysiology experiments and Ms Inna Golyagina for help with analysis of electron microscopy images.

## Author contributions

Y.D., M.D., and O.T. conducted the electrophysiological experiments, N.G. and I.K. analyzed the electron microscopy data, Y.D., N.G., and A.S. performed statistical analysis and prepared the figures, Y.D. and A.S. wrote the manuscript; A.S. designed the experiments; all authors edited the manuscript.

## Data and materials availability

All data needed to evaluate the conclusions in the article are present in the article and/or supplementary materials. Additional data related to this article may be requested from the authors.

