## Supplementary information for "Attenuation of the extracellular matrix increases the number of synapses but suppresses synaptic plasticity"

**Table S1 Full report of two-way RM ANOVA used in the paper**

| Figure panel | Effect of ChABC treatment, $F_{ChABC}$, $p_{ChABC}$ | Effect of stimulation, $F_{stim}$, $p_{stim}$ | Effect of interaction, $F_{ChABC*stim}$, $p_{ChABC*stim}$ | n, sham/ ChABC |
|---|---|---|---|---|
| 2c | F (1, 80) = 17.23, p < 0.001 | F (7, 80) = 48.29, p < 0.001 | F (7, 80) = 0.52, p = 0.817 | 6/6 |
| 3e | F (1, 500) = 183.58, p < 0.001 | F (49, 500) = 0.36, p = 0.999 | F (49,500) = 0.66, p = 0.966 | 6/6 |
| 3f | F (1, 500) = 278.44, p < 0.001 | F (49, 500) = 0.43, p = 0.999 | F (49,500) = 1.05, p = 0.392 | 6/6 |
| 4b | F (1, 100) = 21.99, p < 0.001 | F (9, 100) = 71.67, p < 0.001 | F (9, 100) = 1.72, p = 0.095 | 6/6 |
| 4d | F (1, 100) = 25.96, p < 0.001 | F (9, 100) = 6.18, p < 0.001 | F (9, 100) = 5.41, p < 0.001 | 6/6 |
| 4j | F (1, 100) = 0.01, p = 0.909 | F (9, 100) = 66.76, p < 0.001 | F (9, 100) = 0.08, p = 0.999 | 6/6 |
| 4l | F (1, 108) = 0.41, p = 0.521 | F (5, 108) = 5.34, p < 0.001 | F (5, 108) = 5.35, p < 0.001 | 6/6 |
| 5d | F (1, 600) = 3.17, p = 0.076 | F (49, 600) = 6.87, p < 0.001 | F (49, 600) = 0.29, p = 1.000 | 7/7 |
| 5e | F (1, 600) = 0.97, p = 0.325 | F (49, 600) = 6.52, p < 0.001 | F (49, 600) = 0.12, p = 1.000 | 7/7 |
| 5i | F (1, 600) = 1.81, p = 0.179 | F (49, 600) = 2.20, p < 0.001 | F (49, 600) = 0.11, p = 1.000 | 7/7 |
| 5j | F (1, 600) = 2.93, p = 0.088 | F (49, 600) = 3.06, p < 0.001 | F (49, 600) = 0.46, p = 0.999 | 7/7 |
| S2b | F (1, 40) = 0.36, p = 0.554 | F (3, 40) = 3.43, p = 0.026 | F (3, 40) = 0.15, p = 0.927 | 6/6 |
| S4e | F (1, 300) = 1.22, p =0269 | F (24, 300) = 0.67, p = 0.884 | F (24,300) = 0.46, p = 0.986 | 7/7 |
| S5b | F (1, 80) = 16.75, p < 0.001 | F (9, 80) = 56.56, p < 0.001 | F (9, 80) = 2.034, p = 0.045 | 5/5 |
| S5e | F (1, 80) = 4.89, p = 0.030 | F (9, 80) = 43.88, p < 0.001 | F (9, 80) = 0.68, p = 0.728 | 5/5 |
| S7b | F (8, 72) = 13.40, p < 0.001 | F (8, 72) = 14.14, p < 0.001 | F (8, 72) = 0.16, p = 0.995 | 5/5 |

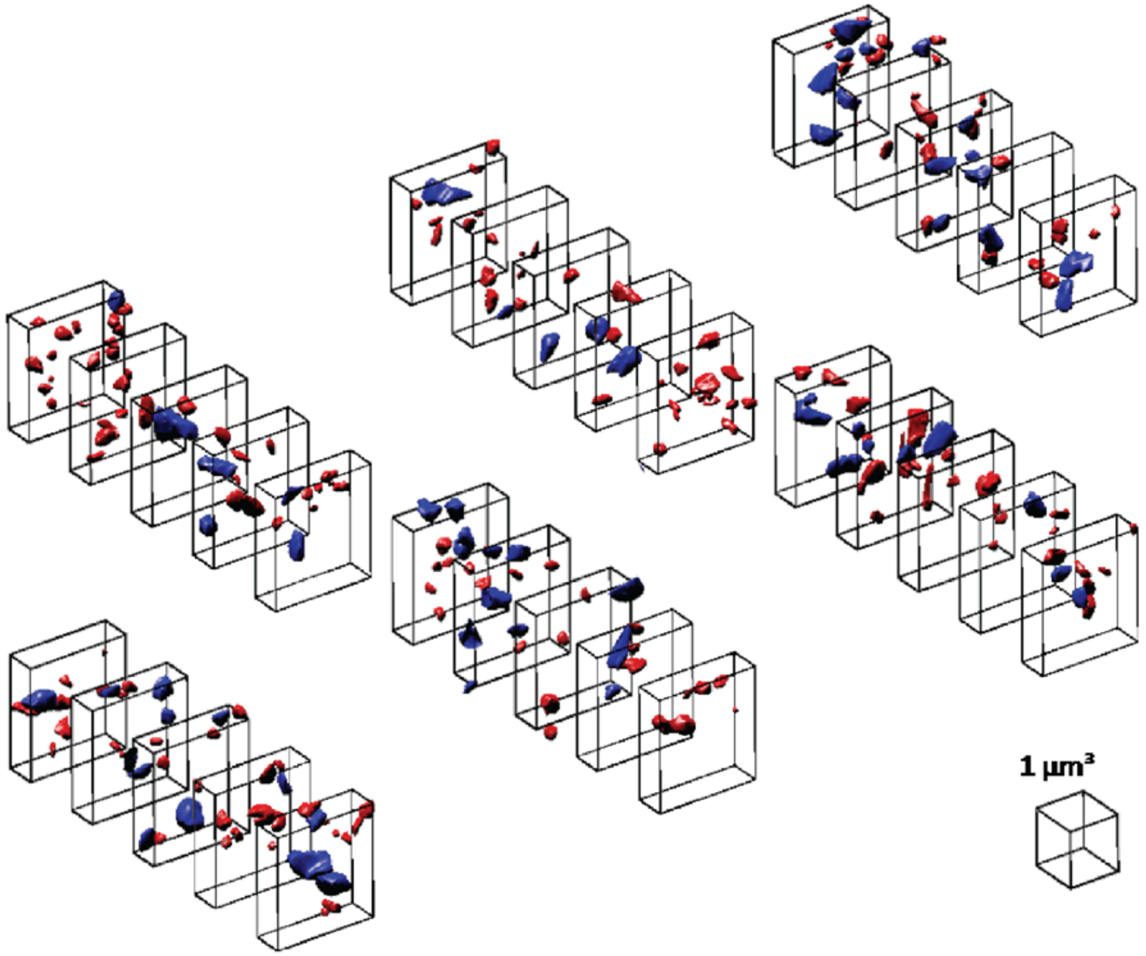


**Figure S1. 3D reconstruction of mushroom and thin spines in one dataset**

To avoid selection bias in the analysis of electron microscopy data, the sampling grid of unbiased bricks was positioned randomly in each image dataset. The datasets were obtained from two animals: sham- and ChABC-treated. Two slices from each animal were used. In total 4 datasets were analyzed.

*Red* – thin spines, *blue* – mushroom spines. The length of a cube side is 1 µm.

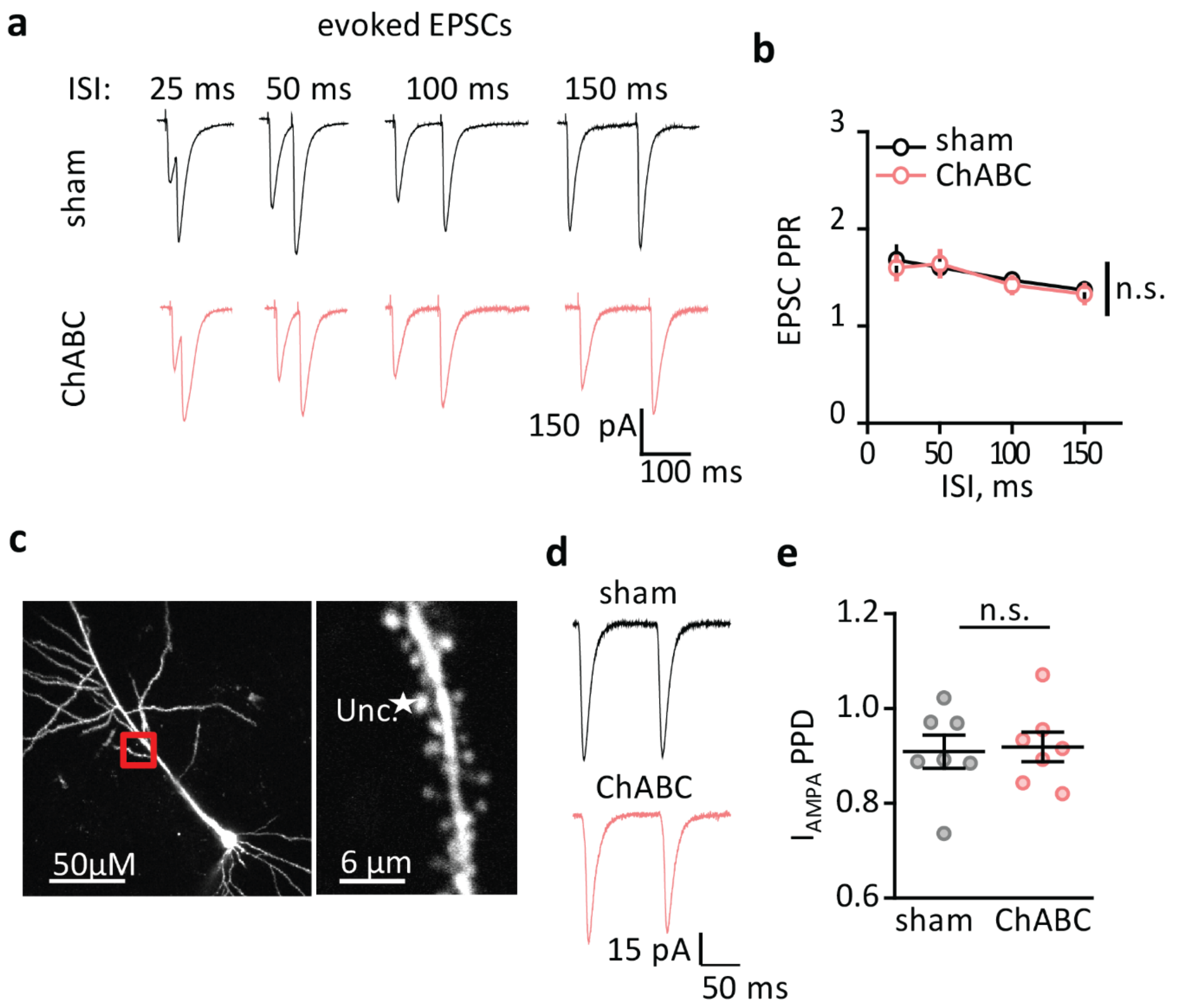


**Figure S2. ECM attenuation does not affect glutamate release probability or lateral diffusion of AMPA receptors in CA1 pyramidal neurons**

**a.** EPSCs evoked by paired-pulse stimulation at different inter-stimulus intervals (ISI, 25 ms, 50 ms, 100 ms, 150 ms). **b.** Summary of the EPSC PPR for different ISI. **c.** *Right panel,* two-photon fluorescent image of CA1 pyramidal neuron loaded through a patch pipette with Alexa 594 (50 µM). The red box represents the area on the dendritic tree where uncaging was performed. *Left panel,* magnification of an oblique dendrite with spines. Star indicates the site of uncaging. **d.** A current ($I_{AMPA}$) recorded in the soma in response to paired-pulse NMI-glutamate uncaging in front of the dendritic spine (panel *c*) in the presence of D-APV (NMDA receptor blocker). **e.** Paired-pulse depression (PPD) of $I_{AMPA}$ due to the desensitization of AMPA receptors by the first uncaging pulse.

*Black traces and circles* – recordings in sham-treated slices; *pink traces and circles* - recordings in ChABC-treated slices. The data are presented as the mean ± SEM. n.s. $p > 0.05$; two-way RM ANOVA (panel *b*); two-tailed two-sample *t*-test (panel *e*).

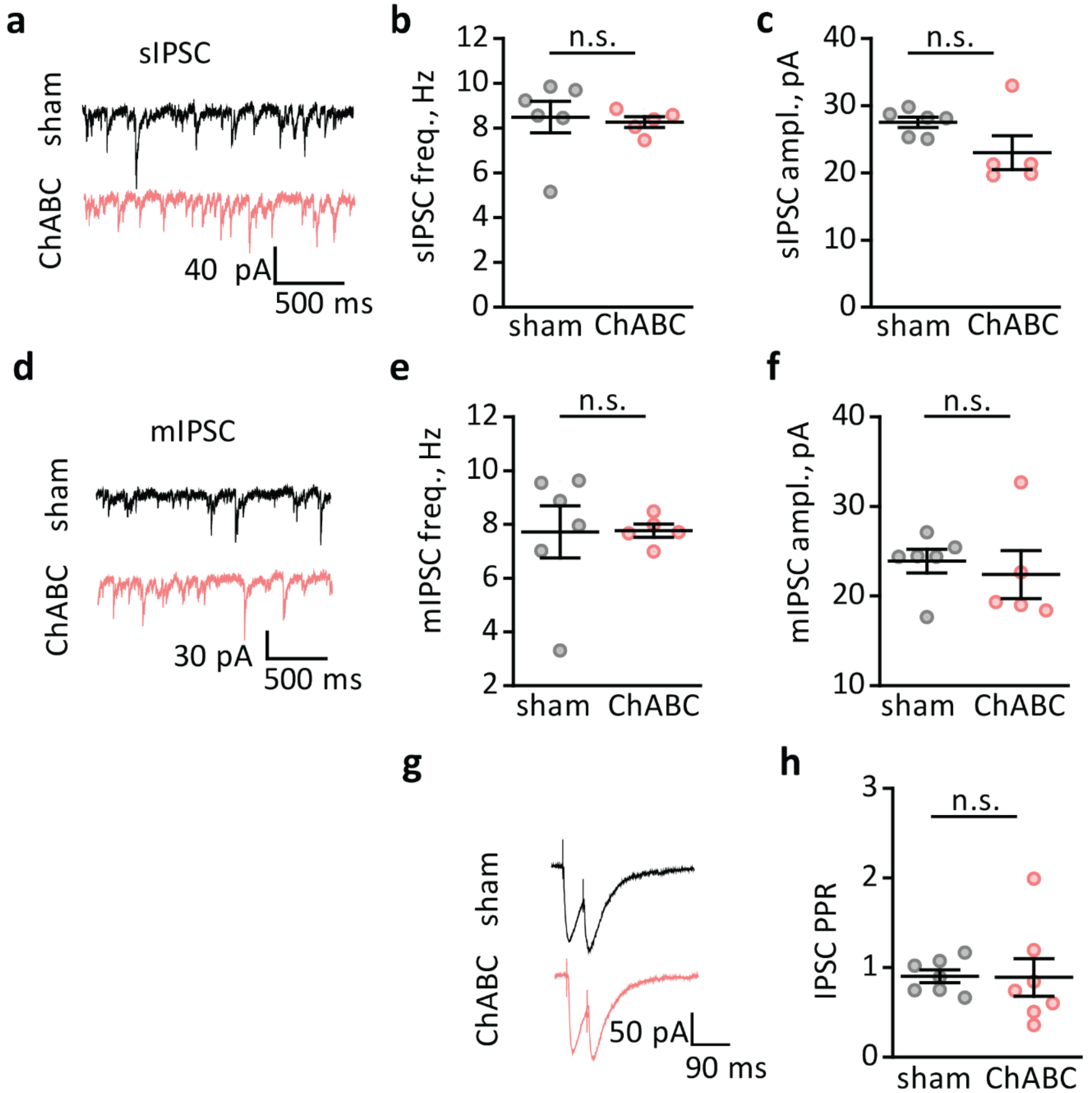


**Figure S3. ECM attenuation does not affect inhibitory transmission**

**a.** Representative traces showing sIPSCs recorded in CA1 pyramidal neurons. **b,c.** Summary of the sIPSC frequency (*b*) and mean amplitude (*c*). **d.** Representative traces showing mIPSCs recorded in CA1 pyramidal neurons in the presence of tetrodotoxin. **e,f.** Summary of the mIPSC frequency (*e*) and mean amplitude (*f*). **g.** IPSCs in response to paired stimulation. **h.** Summary of the paired-pulse ratio (PPR) of IPSCs.

*Black traces and circles* – recordings in sham-treated slices; *pink traces and circles* - recordings in ChABC-treated slices. The data are presented as the mean ± SEM. n.s. $p > 0.05$; two-tailed two-sample *t*-test.

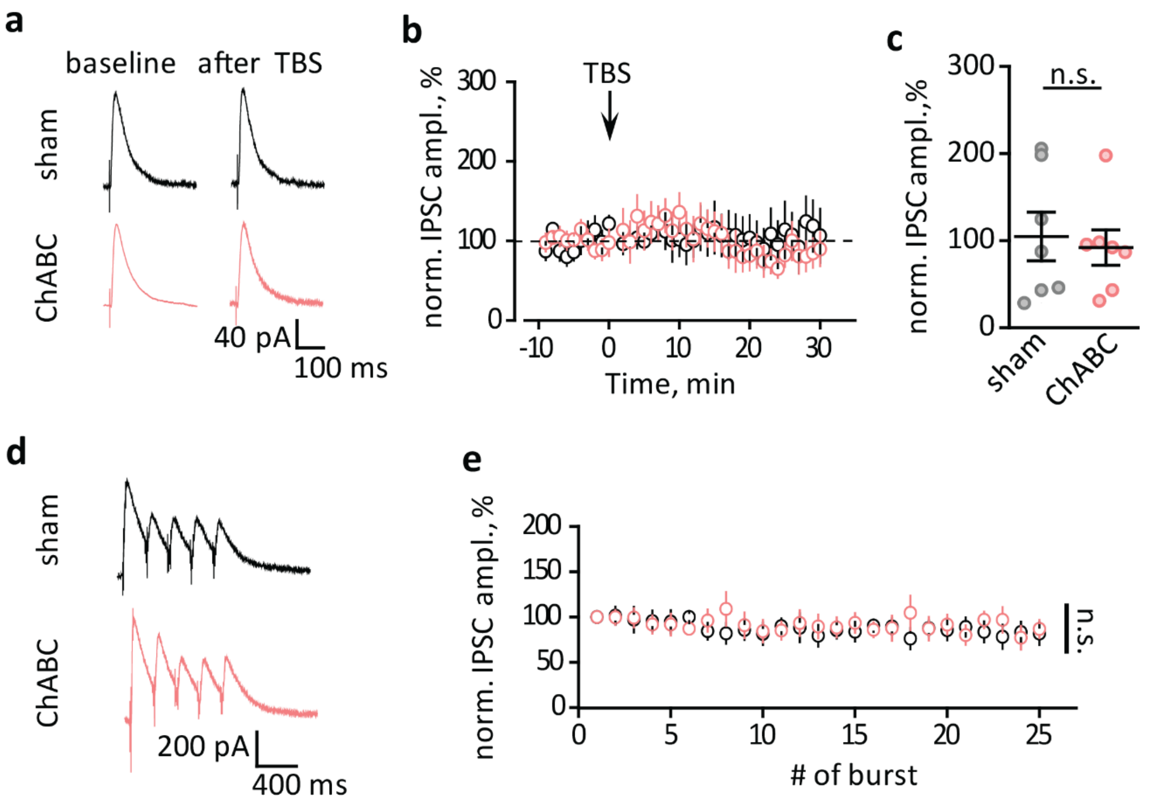


**Figure S4. ECM attenuation does not affect the activity-dependent response of inhibitory synapses**

**a.** IPSCs recorded in CA1 pyramidal neurons before (baseline) and after TBS. AMPA and NMDA receptors were not pharmacologically blocked in these experiments. The neurons were filled with Cs gluconate based intracellular solution and held at a membrane potential of 0 mV, which corresponds to the reversal potential of EPSCs. **b.** The time-course of normalized IPSC amplitudes before and after TBS (arrow, zero time point). **c.** Normalized IPSCs amplitudes averaged over the last 10 min of recordings presented in *b*. **d.** Cell responses to 5 bursts of TBS. **e.** The IPSC amplitude recorded in response to individual bursts during TBS (5 series by 5 bursts).

*Black traces and circles* – recordings in sham-treated slices; *pink traces and circles* - recordings in ChABC-treated slices. The data are presented as the mean ± SEM. n.s. $p > 0.05$; two-tailed two-sample $t$-test (panel $c$); two-way RM ANOVA (panel $e$).

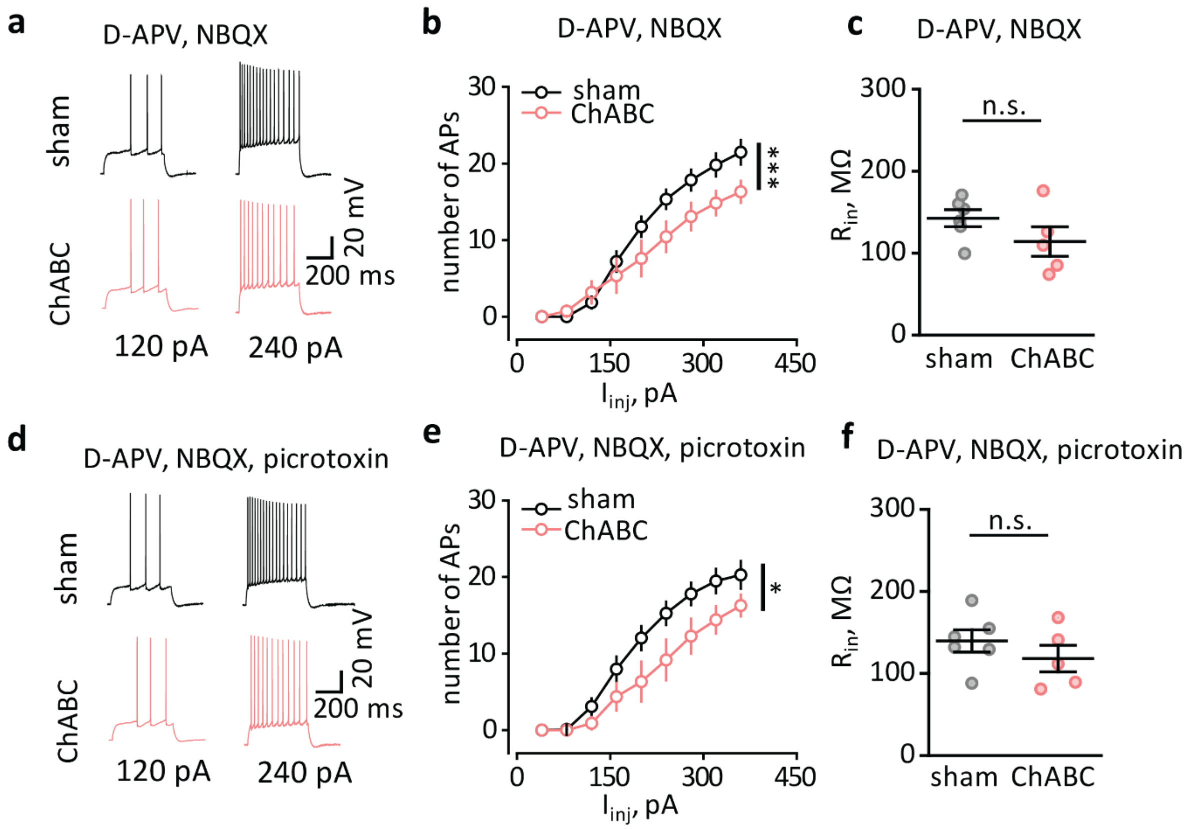


**Figure S5. Decreased excitability of CA1 pyramidal neurons is not mediated by glutamatergic or GABAergic ionotropic receptors**

**a.** Action potentials elicited by depolarizing current steps (120 pA and 240 pA) in the presence of NMDA and AMPA receptor blockers, D-APV and NBQX, respectively. **b.** Summary of the number of action potentials (APs) that occur in response to depolarizing steps in the presence of NMDA and AMPA receptor blockers. **c.** Summary of the input resistance ($R_{in}$) in the presence of NMDA and AMPA receptor blockers. **d.** Action potentials elicited by depolarizing current steps (120 pA and 240 pA) in the presence of NMDA, AMPA and $GABA_A$ receptor blockers, D-APV, NBQX, and picrotoxin, respectively. **e.** Summary of the number of APs that occur in response to depolarizing steps in the presence of NMDA, AMPA, and $GABA_A$ receptor blockers. **f.** Summary of the input resistance ($R_{in}$) in the presence of NMDA, AMPA, and $GABA_A$ receptor blockers.

*Black traces and circles* – recordings in sham-treated slices; *pink traces and circles* - recordings in ChABC-treated slices. The data are presented as the mean ± SEM. n.s. $p > 0.05$; *$p < 0.05$; ***$p < 0.001$; two-way RM ANOVA (panels *b,e*); two-tailed two-sample *t*-test (panels *c,f*).

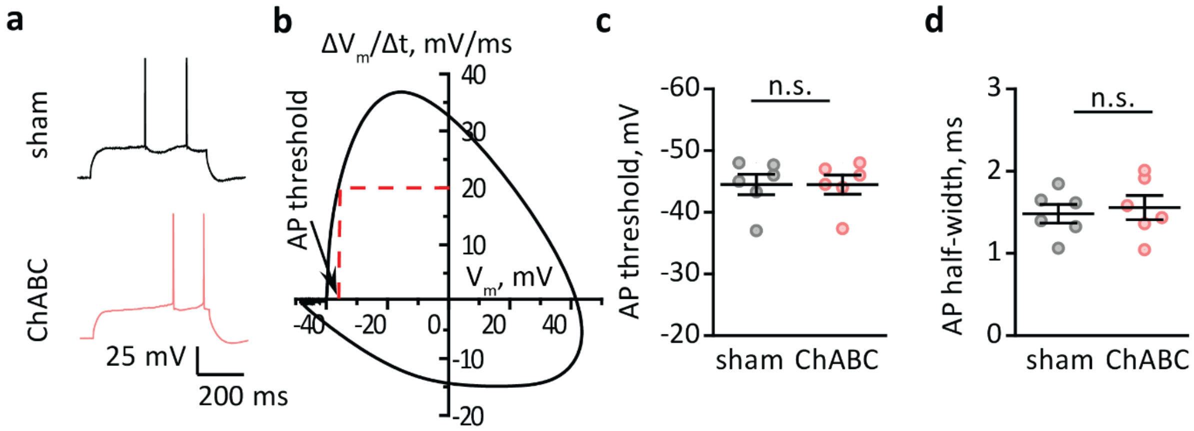


**Figure S6. ECM attenuation does not affect action potential threshold and half-width**

**a.** Action potentials recorded in response to rheobase current injection in CA1 pyramidal neurons. **b.** Phase plane analysis, membrane potential change ($\Delta V_m/\Delta t$) versus membrane potential ($V_m$) for the first action potential generated by the current injection. Action potential (AP) threshold was obtained as $V_m$ at which $\Delta V_m/\Delta t$ = 20 mV/ms (*red dashed line*) was reached. **c,d.** Summary of the AP threshold (*c*) and half-width (the width at half maximum, *d*).

*Black traces and circles* – recordings in sham-treated slices; *pink traces and circles* - recordings in ChABC-treated slices. The data are presented as the mean ± SEM. n.s. $p > 0.05$; two-tailed two-sample *t*-test.

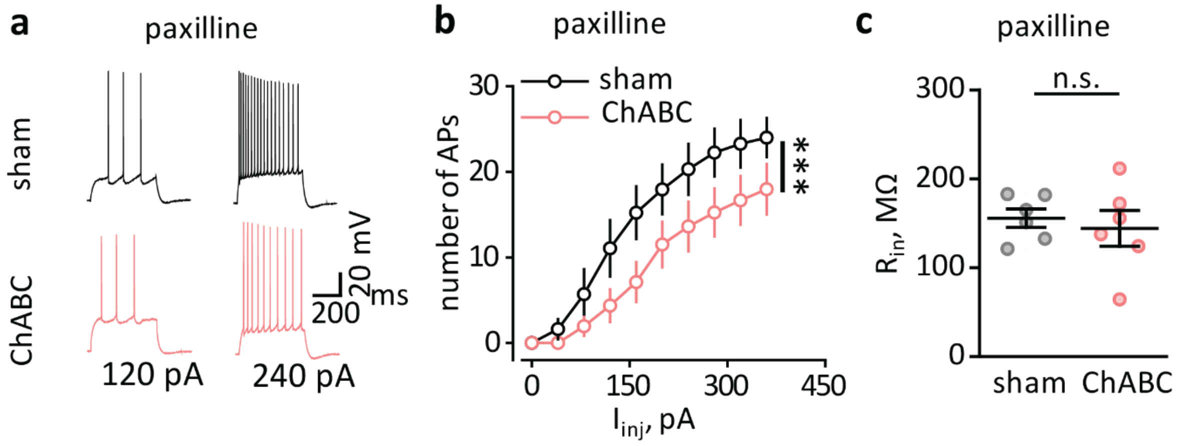


**Figure S7. The reduction of excitability induced by ChABC treatment is not mediated by BK-channels**

**a.** Action potentials (APs) elicited by depolarizing current steps (120 pA and 240 pA) in the presence of the BK channel blocker, paxilline. **b.** Summary of the number of APs that occur in response to depolarizing steps in the presence of the BK channel blocker. **c.** Summary of the input resistance ($R_{in}$) in the presence of the BK channel blocker.

*Black traces and circles* – recordings in sham-treated slices; *pink traces and circles* - recordings in ChABC-treated slices. The data are presented as the mean ± SEM. n.s. $p > 0.05$; ***$p < 0.001$; two-way RM ANOVA (panel *b*); two-tailed two-sample *t*-test (panels *c*).
